# Pulsed-electron illumination does not reduce beam damage for imaging biological macromolecules

**DOI:** 10.1101/2025.07.29.667395

**Authors:** Vishal Kumar, Julika Radecke, K.V. Chinmaya, Inayathulla Mohammed, Ricardo C. Guerrero-Ferreira, Daniel Harder, Dimitrios Fotiadis, Henning Stahlberg

## Abstract

Radiation damage remains a fundamental limitation in cryo-electron microscopy (cryo-EM), constraining the total electron dose that can be used and thus limiting high-resolution imaging of biological specimens. Recent studies have proposed that temporally structured or pulsed electron beams could reduce radiation damage by allowing time for energy dissipation between individual electron interactions. To evaluate this hypothesis, we conducted a systematic investigation using a radiofrequency (RF) driven 300 kV Titan Krios microscope equipped with cold field emission gun (c-FEG) to generate highly regular pulsed electron beams for specimens under cryogenic conditions. We compared radiation damage in three representative samples: paraffin 2D crystals, bacteriorhodopsin (purple membrane) 2D crystals, and plunge-frozen tobacco mosaic virus (TMV) in vitreous ice, under both pulsed and conventional (= random) illumination, while keeping all other imaging conditions constant. Radiation damage was quantified by tracking the decay of computed diffraction intensities to determine the critical dose (*N*_*e*_). We observed no statistically significant difference in critical dose between pulsed and random illumination across all 3 samples. Our findings provide a critical reference point for future development and evaluation of temporally modulated electron sources in cryo-EM instrumentation.

## Introduction

Cryo-electron microscopy (Cryo-EM) has revolutionized structural biology by enabling the visualization of biological macromolecules in three dimensions (3D) at near-atomic resolution.^1,2^ This breakthrough has been driven by recent advancements in electron microscopy hardware, direct electron detectors, automated data acquisition strategies, and significant improvements in image processing algorithms that are available in user-friendly data analysis software packages.^1–7^ Together, these innovations have empowered researchers to investigate complex biological processes at the molecular level, opening new avenues for discovery in fundamental biology, drug development, soft functional materials, and beyond.

Despite significant advances in instrumentation and data processing, radiation damage remains a fundamental barrier to further improve the performance of cryo-electron microscopy (cryo-EM).^8,9^ It is well established that radiation from the electron beam disrupts molecular bonds within biological macromolecules, leading to structural degradation and conformational changes under continuous exposure.^10^ While the precise mechanisms and contributing factors of radiation damage remain active areas of investigation, several practical strategies have proven effective in mitigating its effects. The most important measure to reduce the consequences of beam damage is cooling specimens to cryogenic temperatures, typically liquid nitrogen (∼77 K)^11^ or liquid helium (∼4 K).^12–15^ This significantly reduces the mobility of ionized species, thereby limiting the extent of radiation-induced damage. The use of higher accelerating voltages (e.g., 300 keV versus 100 keV) has also been shown to reduce radiation damage as the inelastic scattering cross-section reduces at higher accelerating voltage.^16^ However, the elastic scattering cross-section drops even more steeply at higher accelerating voltages. As a result, 100 keV is preferable for thinner samples, whereas 300 keV is advantageous for thicker specimens, such as cellular sections used in cryo-electron tomography.^16^

Recently, several groups have begun to debate whether radiolysis is influenced by the temporal profile of electron exposure. For example, Flannigan et al. reported that a pulsed electron beam can reduce radiolysis-induced damage by a factor of two in paraffin.^17^ Their hypothesis was that the time between electron pulses allows sufficient recovery for the sample to dissipate reactive atom species at the local excitation site. However, their study was limited to very low cumulative doses (<0.1 e^−^/Å^2^) due to the long acquisition times required by their laser-excited ultrafast electron beam setup. Their work therefore was unable to probe the beam damage effect at doses relevant for cryo-EM.

In another study, Kisielowski et al. reported an 80x to 100x reduction in damage when using pulsed electron beams on MgCl_2_.^18^ However, their experimental conditions for pulsed and random illumination were strongly different: The local brightness was orders of magnitude higher in random vs. pulsed illumination mode, raising questions about the comparability of the results. Since sample temperature does affect the consequence of beam damage and phase transition, the much higher brightness in non-pulsed operation might have created a locally very high temperatures, which could explain the observed differences in transition speed of MgCl_2_ to another crystal form.

In yet another study, Choe et al. observed a two-fold increase in the critical dose, defined as the dose at which the intensity of diffraction spots falls to 1/e of the original, using a pulsed beam configuration.^19^ They collected their data at extremely low dose rates (∼0.02 e^−^/Å^2^s), which limits its applicability to practical data acquisition strategies. That work did not probe dose values relevant for cryo-EM, which require 50 e^−^/Å^2^ or more. Recording such data at that dose rate would result in ∼ 40 minutes of data collection per micrograph, which is not feasible if many thousands of micrographs are required.

In contrast to these findings, Zhao et al. recently reported no improvement in the critical dose when applying pulsed electron beams to paraffin, using a laser-driven photocathode system.^20^ However, such modified photocathode setups are less suitable for automated, high-resolution cryo-EM due to their broader energy spread and lower brightness compared to conventional cold field emission guns (c-FEGs). Moreover, paraffin is a relatively simple hydrocarbon, while proteins consist of complex assemblies of amino acids with diverse structural and chemical properties, and these are embedded in vitreous water in cryo-EM studies. These differences may significantly influence the dynamics of radiation damage under pulsed electron illumination for cryo-EM specimens.

The possibility of reducing the impact of electron beam radiation damage would represent a massive improvement for cryo-EM. Increasing the allowed electron dose for high-resolution imaging of biological macromolecules from the currently possible 100 e^−^/Å^2^ to 200 e^−^/Å^2^ would be equivalent to an increase in the detection quantum efficiency (DQE) of the camera by a factor of two, which would have radical improvements in the performance of cryo-EM as consequence.^21^ It would for example significantly lower the size of the smallest possible protein particle that can be imaged by cryo-EM,^22^ or it would allow a much finer classification of proteins or protein conformations in cryo-EM of single particles. It would also give significant improvements in the possibilities for detecting and identifying even smaller protein particles in cryo-electron tomography reconstructions of biological tissue. Considering the importance of improving the signal-to-noise ratio (SNR) of cryo-EM imaging, systematic investigations of the impact of pulsed electron beam illumination for cryo-EM specimens under relevant conditions are important.

Here, we present a systematic analysis of the potential of pulsed electron beam illumination for imaging biological specimens at cryogenic temperatures. We employed a radiofrequency (RF) driven c-FEG system mounted on a 300kV high-end Titan Krios electron microscope to generate highly repetitive pulsed electron illumination for imaging biological macromolecules at cryo-temperatures. Radiation damage was assessed by monitoring the fading of computed diffraction intensity across all samples, from which the critical electron dose (*N*_*e*_) values were calculated. We present comparisons of electron beam damage between pulsed and random (conventional) electron beam configurations under otherwise identical imaging conditions, including identical electron dose rates and identical exposure times. We conducted imaging on three different specimens: paraffin 2D crystals (C36H74) on continuous carbon film, bacteriorhodopsin 2D membrane crystals (purple membrane) embedded in glucose, and plunge-frozen tobacco mosaic virus (TMV) embedded in vitreous ice, all at liquid-nitrogen cryo temperatures and using both random and pulsed electron beam illumination. Under comparable exposure conditions, our results show no significant differences in damage profiles and *N*_*e*_ across all three samples, indicating that pulsed electron beam illumination provides no substantial benefit over random electron beam exposure for imaging biological samples at cryogenic temperatures.

## Results and Discussion

Pulsed electron beams can be generated using a photocathode system, where a high-energy laser pulse induces the emission of electron bunches via the photoelectric effect.^23,24^ These electrons are then accelerated by high-voltage electric fields and subsequently focused using electromagnetic lenses to form pulsed electron beam. However, such modified photocathode systems often suffer from a large energy spread and low brightness of the produced electron beam, making them unsuitable for high-resolution and automated data acquisition. Modern electron microscopes are increasingly equipped with cold Field Emission Guns (c-FEGs), which offer a very low energy spread (∼0.3 eV) and high brightness. These characteristics played a pivotal role in pushing the resolution limits, particularly in cryo-electron microscopy (cryo-EM) applications, allowing resolutions of up to 1.09 Å for single protein particles.^25^ For the work presented here, a c-FEG–equipped 300kV Titan Krios microscope was equipped with a dual-mode RF cavity system developed by DrX.Works,^26^ enabling the generation of pulsed electron beam illumination.

**Figure 1a** illustrates the mechanism of pulsed electron beam generation, where a continuous electron beam is focused into a dual-mode RF cavity. This cavity, driven at 2.416 GHz and 2.4915 GHz, generates a rapidly oscillating electromagnetic field that deflects the beam in a transverse Lissajous pattern (**Figure 1b**). To produce a pulsed electron beam, a 50 µm C2 aperture was inserted downstream, which chopped the beam and allowed only the desired portion of the beam (only individual segment rather than 4 segments created by dual-mode cavity) to pass through. It should be noted that the individual segments from the Lissajous pattern are rectangular, while we used a round aperture. To mitigate this, we carefully adjusted the spot size, magnification and condenser lens settings, so that the electron beam fully covered the aperture, resulting in a homogeneous pulsed electron beam. The temporal profile of this beam is shown in **Figure 1c**, demonstrating that no electron events were able to occur between the respective electron pulses.

**Fig. 1.**
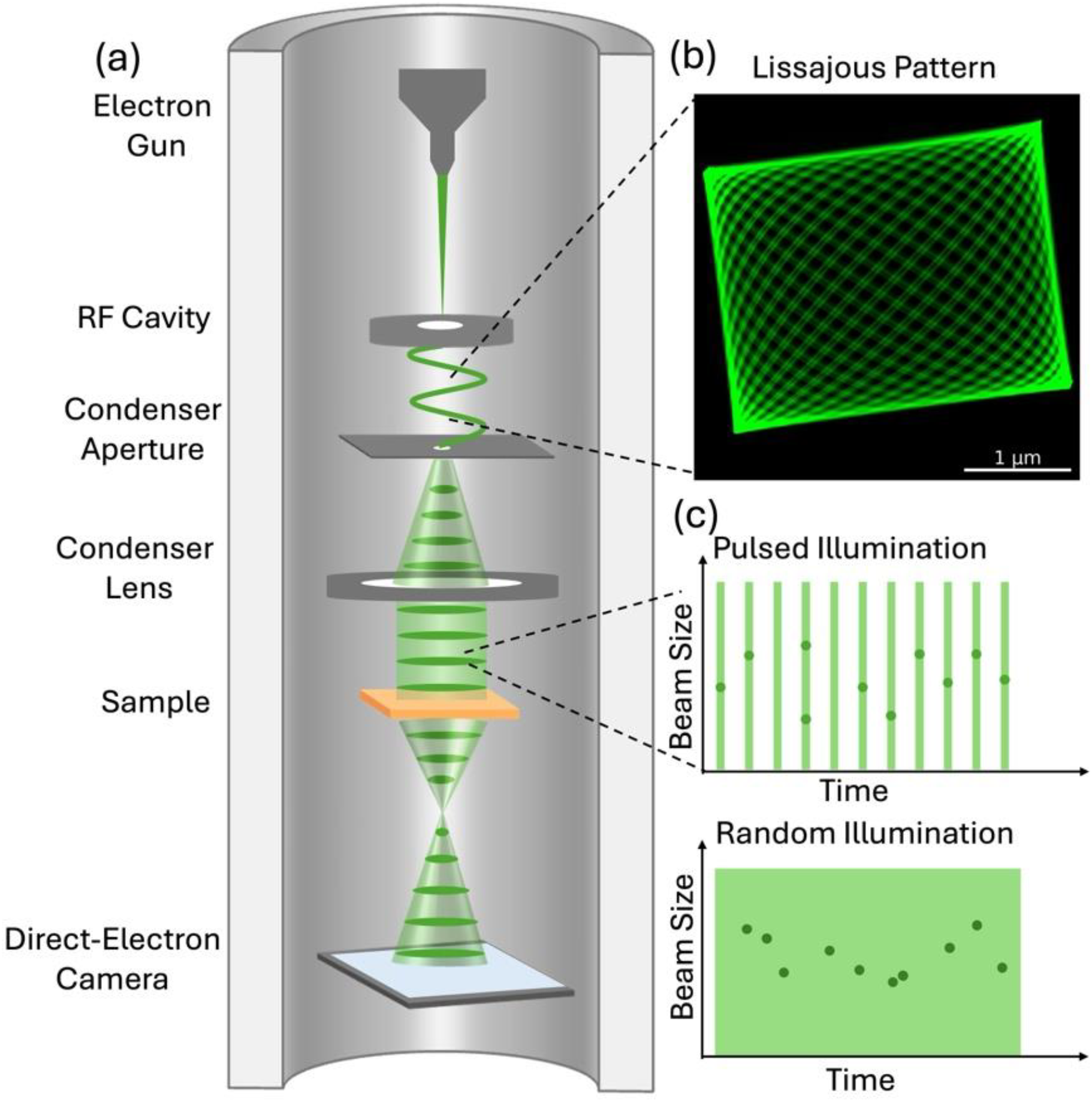
The experimental setup for pulsed electron illumination. **a** Illustration of a c-FEG equipped 300kV Titan Krios electron microscope, modified by a dual mode RF cavity system for pulsed electron beam generation. **b** A slightly elliptic cavity drives beam deflection in X and Y at 2.416 GHz and 2.4915 GHz respectively, after which the beam forms a Lissajous pattern (experimental image). A pulsed electron beam is formed by using a small condenser aperture to cover only a part of the Lissajous pattern. **c**. Temporal profile for the pulsed and random illumination configuration, with individual pulses has spacing of 13.33 ns. (Note: this is just a schematic, the actual electron counts within individual pulses can vary according to experimental settings)

In our experiments, the resulting electron pulse bunch width ranged from 0.9 to 1.1 picoseconds, and the pulse repetition rate was 75 MHz (corresponding to a 13.33 ns gap between individual pulses), determined by the frequency difference between the two RF modes. The resulting pulsed electron beam exhibited uniform illumination (**Supplementary Figure 2**), maintained long-term intensity stability (**Supplementary Figure 3**), and did not hinder high-resolution imaging, as demonstrated using a gold grating sample (**Supplementary Figure 4 and 5**). It is important to note that under typical experimental conditions, approximately 75% of the electron bunches were empty, while only about 25% contained single-electron events, as detailed in **Supplementary Table 1**. This ensured that, on average, only one electron arrived at the sample at any time, minimizing multi-electron scattering events. For comparison, the temporal profile of a conventional electron beam is also shown, which was obtained by switching off the RF cavity and adjusting the condenser lens settings so that the same beam brightness and diameter was achieved. In this mode, due to the absence of temporal gating, the time interval between individual scattering events cannot be determined. Therefore, we refer to this configuration as the random-in-time (random or conventional) illumination mode.

To evaluate the influence of pulsed versus random illumination, we performed real-space imaging on three different samples. Imaging parameters such as electron dose rates, defocus values, and exposure times were kept the same to eliminate any additional effects. By tracking the intensity decay of the computed diffraction spots in reciprocal space, we determined the radiation damage profile. From this, we calculated the critical electron dose (*N*_*e*_) at which the spot intensity had fallen below 1/e of the initial value. This was used to determine the impact of radiation damage using both configurations.

**Figure 2a** shows a representative power spectrum of paraffin crystals, clearly displaying intensity spots corresponding to the (1,1) plane at 4.12 Å and the (2,0) plane at 3.71 Å, along with their respective Friedel pairs. The mean intensity of all six computed diffraction spots is calculated and referred to as the 4 Å resolution band. **Supplementary Figure 6** shows the fading of spot intensities with accumulated electron dose over time. The radiation damage profiles, obtained from 10 independent measurements under both pulsed and random illumination conditions, are shown in **Figure 2b**. These profiles indicate no significant advantage of pulsed illumination over random illumination in mitigating radiation damage. This conclusion is further supported by comparable *N*_*e*_ values (mean values: 11.15 ± 1.47 e^−^/Å^2^ for pulsed and 11.71 ± 0.94 e^−^/Å^2^ for random illumination, all values tabulated in **Supplementary Table 2**) observed for both modes, as shown in **Figure 2c**. *N*_*e*_ values were calculated by fitting a single-exponential decay curves to the damage profiles and determining the doses at which the intensities fall below the 1/e threshold.

**Fig. 2.**
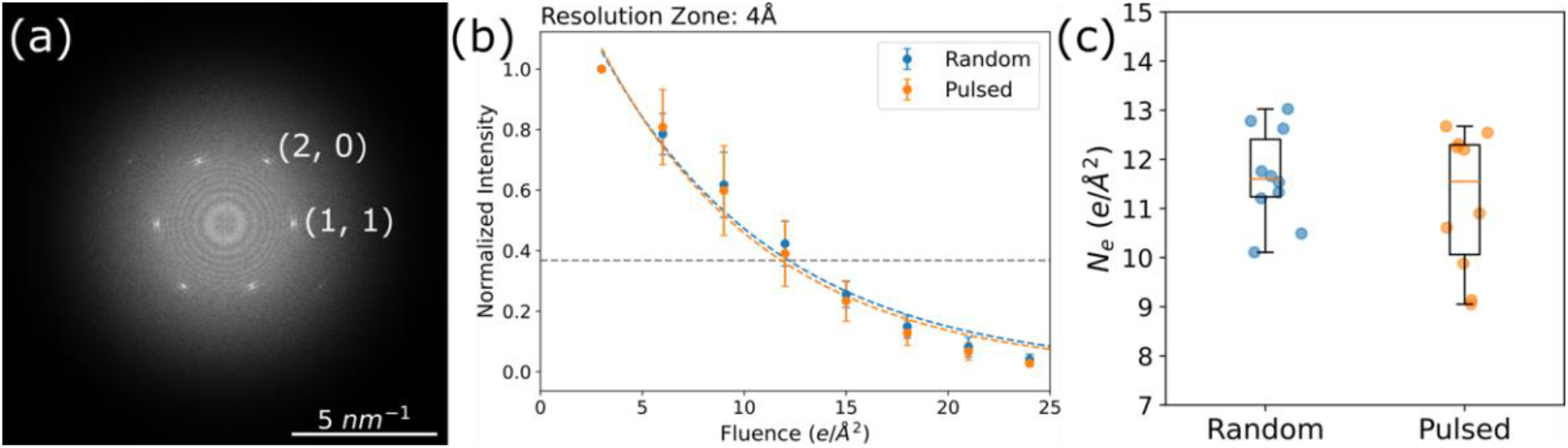
Electron beam damage measured by imaging paraffin crystals. **a** Representative computed power spectrum of paraffin crystal image, **b** normalized spot intensities with accumulated electron fluence in the 4 Å resolution band (from (1,1) and (2,0) planes) and corresponding exponential fits for random and pulsed electron-beam illumination marked by dashed lines. Error bars represent the standard deviation of the measurements. The horizontal line represents a 1/e threshold for calculating *N*_*e*_ values. **c** *N*_*e*_ dose values for random and pulsed illumination.

For protein samples, we first tested purple membrane crystals, embedded in glucose on continuous carbon film, and then cooled to LN_2_ temperature inside the electron microscope. The purple membranes are composed of a phospholipid bilayer with hexagonally arranged 2D crystals of trimers of the protein bacteriorhodopsin.^27^ The crystals in p3 symmetry have a large unit cell (side length 63 Å), resulting in numerous computed diffraction peaks in reciprocal space, as shown in **Figure 3a**. To monitor radiation damage across different resolution ranges, we divided the Fourier space into multiple resolution zones:

**Fig. 3.**
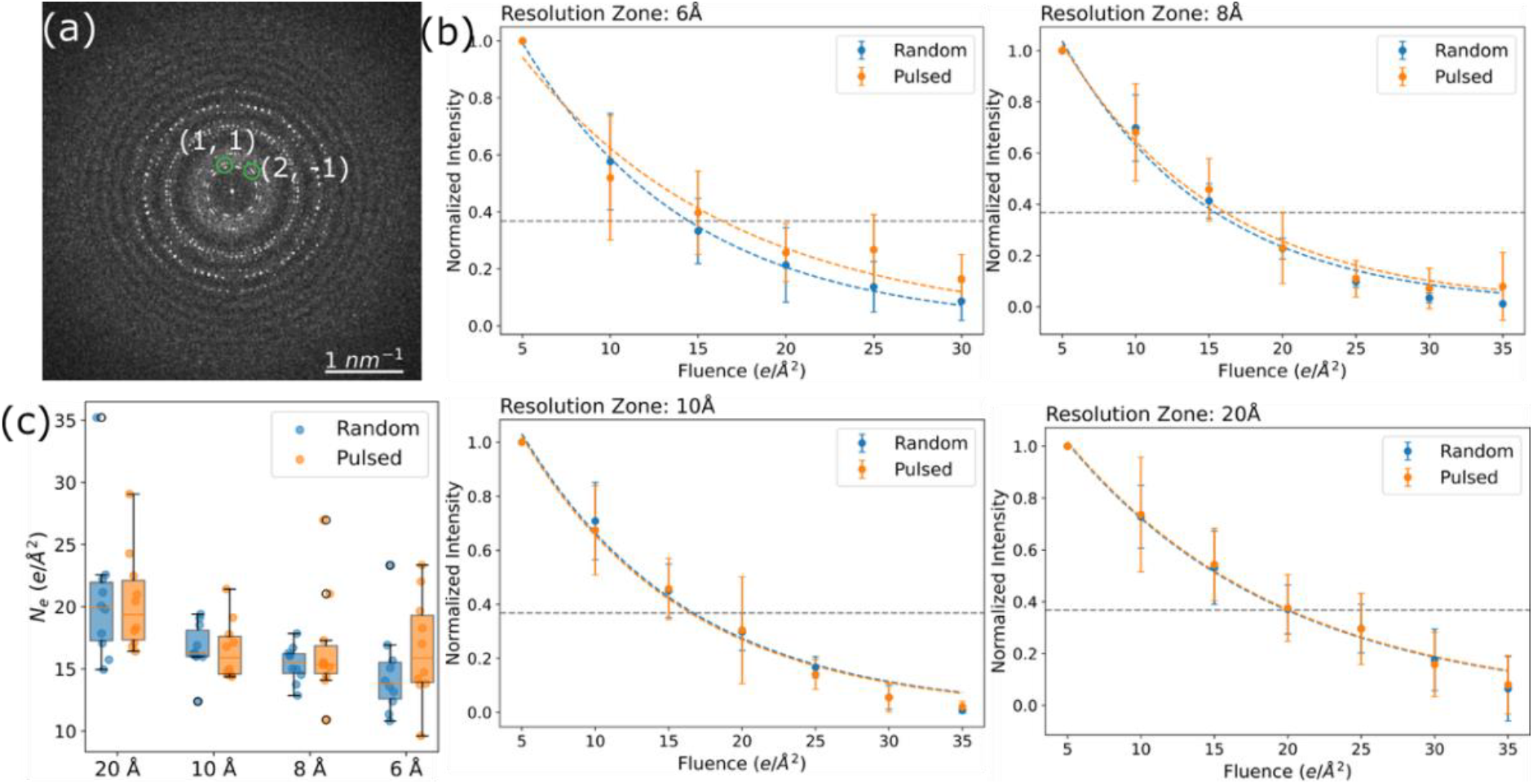
Electron beam damage measured by imaging purple membrane 2D crystals. **a** Representative power spectrum of purple membrane crystal image, with planes (1,1) and (2, -1) marked on the image. **b** Normalized spot intensities with accumulated electron fluence in different resolution bands (20 Å: 12-35 Å, 10 Å: 9-12 Å, 8 Å: 7-9 Å, 6 Å: 6-7 Å) and corresponding exponential fits for random and pulsed electron-beam illumination marked by dashed lines. Error bars represent the standard deviation of the measurements. The dotted horizontal lines represent 1/e thresholds for calculating *N*_*e*_ values. **c** *N*_*e*_ dose values for random and pulsed illumination in different resolution zones.

- 20 Å zone: 12-35 Å
- 10 Å zone: 9-12 Å
- 8 Å zone: 7-9 Å
- 6 Å zone: 6-7 Å

The progressive fading of spot intensities as a function of accumulated electron dose over time is shown in **Supplementary Figure 8**. The intensity decay within each resolution band was tracked and plotted, as shown in **Figure 3b**. The damage profiles across all resolution zones show no significant difference between pulsed and random illumination. Additionally, *N*_*e*_ values calculated for each resolution zone are also plotted (all values are tabulated in **Supplementary Table 3**) and show no measurable benefit from using pulsed illumination.

Finally, we tested our hypothesis on TMV embedded in vitreous ice, as biological specimens are typically imaged using amorphous ice as the supporting medium. **Figure 4a** shows a representative power spectrum acquired at a low dose (5 e^−^/Å^2^), revealing two characteristic reflections: one at 23 Å and another at 11.5 Å, corresponding to the structural features of TMV.^28^ For our analysis, we focused on the 23 Å peak, as the signal-to-noise ratio (SNR) of the 11.5 Å peak was too low for reliable quantification. Computed diffraction intensities fading with increasing accumulated electron dose over time are shown in **Supplementary Figure 10**. The radiation damage profile for TMV, shown in **Figure 4b**, once again demonstrates no significant difference between pulsed and random illumination. Correspondingly, the mean *N*_*e*_ values or 38.15 ± 3.84 e^−^/Å^2^ for pulsed and 35.22 ± 3.70 e^−^/Å^2^ (all values are tabulated in **Supplementary Table 2**) for random beam illumination, further confirm this trend. In addition, we have collected a small dataset for Single Particle Analysis (SPA). The data was processed in Relion 5^29^ and the resulting map and FSC curves has been shown in **Supplementary Figure 12**. Additionally, per-frame B-factor plots^30^ (**Supplementary Figure 13**) show similar decay profiles for datasets taken using pulsed and random illumination. Thus, these further strengthen our argument, that a pulsed electron beam doesn’t provide significant improvement over conventional (=random) illumination scheme for imaging biological macromolecules.

**Fig. 4.**
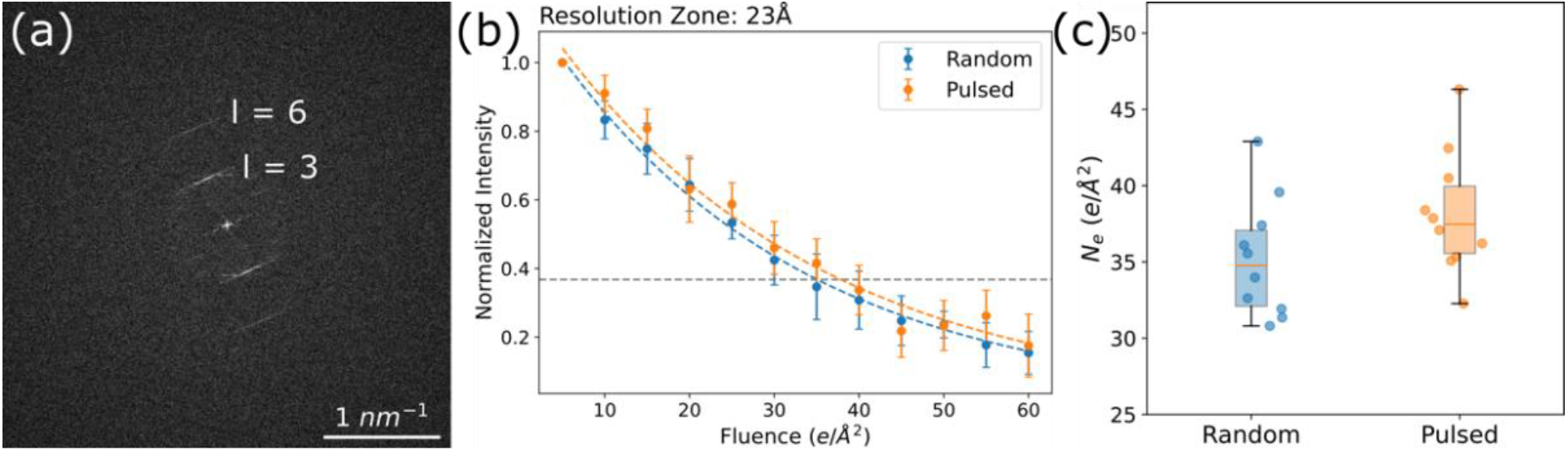
Electron beam damage measured by imaging tobacco mosaic viruses (TMV). **a** Representative power spectrum of TMV embedded in vitreous ice, showing l=3 (23 Å) and l=6 (11.5 Å) layer lines. **b** Normalized spot intensities with accumulated electron fluence in 23 Å resolution band and corresponding exponential fits for random and pulsed electron-beam illumination, marked by dashed lines. Error bars represent the standard deviation of the measurements. The dotted horizontal line represents the 1/e threshold for calculating *N*_*e*_ values. **c** *N*_*e*_ dose values for random and pulsed illumination.

## Discussion

In this study, we investigated the prospect of pulsed electron beam illumination to reduce radiation damage for imaging biological specimens in cryo-electron microscopy (cryo-EM). We used a dual mode RF cavity behind a cold field emission gun (c-FEG) system integrated into a Titan Krios microscope to generate pulsed electron beams, and we compared the effect of pulsed illumination on radiation damage with that of random (conventional) electron illumination. Radiation damage was assessed through fading of computed diffraction intensities, from which the critical dose (*N*_*e*_) was calculated for three representative sample types: paraffin crystals, purple membrane protein crystals, and tobacco mosaic virus (TMV) embedded in vitreous ice.

Across all three systems and different resolution zones, we found no statistically significant difference in radiation damage profiles or critical dose values between pulsed and random illumination modes under equivalent imaging conditions. Flannigan et al. proposed that the time between incoming electrons would allow samples sufficient time to dissipate any localized energy buildup, even though their work only probed very small accumulated electron doses around 0.1 e^−^/Å^2^.^17^ In our study, the interval between consecutive pulses was 13.33 ns, substantially longer than the 160 ps used by Kisielowski et al.,^18^ or the 192 ps used by Choe et al.^19^ Despite the longer time delay between consecutive pulses, we still did not observe any significant improvement. Our results suggest that, at least under the current implementation conditions and dose rates relevant for high-resolution cryo-EM, pulsed electron beam illumination offers no substantial advantage over conventional random illumination in terms of radiation damage mitigation. It can be due to different reasons:

1. We used the time interval of 13.33 ns between the pulses, which is a hardware limit that is imposed by our instrumentation. These 13.33 nanoseconds may still not be sufficiently long for damage recovery. For example, for generated atom radicals to diffuse, diffusion is proportional to the square root of time,^31^

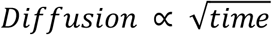 It might be that longer time intervals might be needed between the pulses for effective damage mitigation from radicals.
2. We use a parallel electron beam with an illumination diameter of about 400 nm, which is typically used in normal cryo-EM experiments. Under these conditions, subsequent electrons may land far apart from one another on the specimen, so that a single electron interaction would be spatially isolated, and a subsequent electron can strike well outside that local region. This possibility of spatial separation on the sample plane could lower the likelihood of cascading damage effects. Consistent with this perspective, Kisielowski et al.^32^ recently claimed that a pulsed electron beam was more effective in reducing beam damage when using a smaller irradiation area. In the smaller areas used by Kisielowski et al., the dissipation of local heating and build-up of radicals might require the time interval between incoming electrons (the pulsed electron beam), as electrons arrive very close to each other spatially. In contrast, for the large illumination areas typically used in cryo-EM and also used in our work, we did not observe any measurable benefit from the pulsed beam. This could then be interpreted as that if electrons are already sufficiently separated spatially, then local radicals or thermal effects can relax between individual events even in conventional electron illumination, so that any benefit of pulsed illumination is not required. This observation is consistent with perspectives from Robert Glaeser^8^ and Christopher Russo et. al.,^33^ who proposed that only a small number of electrons are present in the TEM column at any given moment, and that they arrive at the specimen with sufficient spatial separation to produce the same beneficial effect as using pulsed-electron beam.

Given the growing interest in temporally structured electron beams and the considerable technical complexity involved in implementing pulsed systems for cryo-EM, our results serve as a critical reference point. We showed that the reported benefits of pulsed illumination do not apply in the here probed context of structural biology cryo-EM. Further investigations may be warranted at higher temporal resolution, or different pulse repetition rates, but our results caution against widespread adoption of pulsed beam technology without demonstrable performance gains in practical cryo-EM workflows.

## Methods

### Specimen preparation

Paraffin crystals (C36H74) were made by drop casting 2 µL of saturated paraffin solution in hexane (purchased from Sigma-Aldrich) onto amorphous carbon film coated grids. The excess hexane solution evaporated quickly, after which grids were loaded directly into the cryo-stage of the Titan Krios. Purple membrane crystals were synthesized by the method described in previous study.^34^ Carbon-coated grids were first glow-discharged for 40 sec, after which 1µL of purple membrane solution was dispensed onto the grid, followed by 1µL of 2% glucose solution. Grids were blotted and dried in air. Following this, the grids were loaded directly into the cryo-stage of the Titan Krios. TMV samples were prepared for cryo-EM by using Quantifoil 1.2/1.3 grids, which were plasma cleaned. Then 5 nL of TMV solution at 20 mg/ml were deposited on the grids and plunge frozen using a cryoWriter instrument.^35^

### Cryo-electron microscopy data collection

All data were collected with a Titan Krios G4 (Thermo Fisher Scientific, TFS), equipped with a cold-FEG and a Falcon 4i direct electron detector. The microscope has been equipped with a dual-mode RF cavity system from DrX.Works for the generation of ultrafast electron pulses.^26^ For operation, the electron beam was focused into the center of the RF cavity, where the resonant cavity applied strong oscillating electromagnetic fields in X and Y directions to the electron beam, which was deflected laterally to form a Lissajous pattern. The RF Cavity was nominally driven at the resonant frequencies of 2.416 GHz and 2.4915 GHz respectively, as reported by the software user interface. An exact, fixed ratio of 32:33 between the two modes forming a Lissajous pattern was established, resulting in 32 × 33 crossings. A small phase shift between the signal of the two frequencies was introduced to separate out the two beams riding on top of each other, so that a closed loop pattern was formed, along which the beam was scanning in only one direction (**Supplementary Figure 1**). Following this, a C2 aperture (50 µm) was inserted below the cavity, and by adjusting spot size, magnification and illumination intensity, a homogenous electron beam illumination over the surface of the C2 aperture hole was created (**Supplementary Figure 2**). The resulting electron beam had a repetition rate of 75 MHz. Since this operation mode eliminated most of the electron beam to form the short pulses, we used a very high gun emission current of ∼60 nA instead of the normally used ∼5 nA, so that the resulting beam allowed cryo-EM imaging at relatively high dose-rates (∼1 e^-^/Å^2^s). As mentioned in **Supplementary Table 1**, almost 75% of pulses are empty. The number of electrons per pulse is dependent on the illumination current before the RF cavity (function of tip current, aperture size, and spot number) and RF modulation parameters, as a higher magnetic field in cavity (or higher power in both cavity antennae) results in higher amplitude of the beam deviations, thereby shorter traversal times of the beam over the C2 aperture, which creates shorter pulses, leading to less electrons per pulse for a given illumination current.

Beam stability was measured over an extended period of time (**Supplementary Figure 3**). We found no considerable change in current intensity over a period of 60 mins. The current in the pulsed electron beam was measured using a Faraday cup. The RF cavity induced a strong beam tilt, which was compensated for by inducing strong beam tilt from the gun in the opposing direction before the RF cavity. A standard gold grating sample was used for alignment of the system, and to assess the information transfer in pulsed beam operation (**Supplementary Figure 4 and 5**). Conventional random illumination cryo-EM images were collected with the RF cavity switched off, and at conditions that exactly matched the same dose rate, illumination diameter on the sample, and exposure times, all done on the same instrument as the pulsed beam experiments.

All datasets were acquired at liquid nitrogen temperature at 300 keV using a Falcon 4i camera. For data acquisition, paraffin and TMV datasets were acquired at a physical pixel size of 0.65 Å and purple membrane data at a physical pixel size of 0.52 Å. Paraffin data were acquired at a dose rate of 1.35 e^-^/Å^2^s, purple membrane at 2.08 e^-^/Å^2^s, and TMV data at 1.1 e^-^ /Å^2^s. To measure radiation damage, multiple consecutive images were acquired at each specimen location for each dataset using SerialEM.^5^ For paraffin, each image had a dose of 3 e^-^/Å^2^, for purple membrane and TMV each image had a dose of 5 e^-^/Å^2^ to ensure enough SNR in the computed diffraction spots. A small dataset for a single particle analysis with TMV was also acquired at the same settings, with each micrograph having a total dose of 50 e^-^/Å^2^. Both pulsed and random beam experiments were done on the same microscope, using the same specimen grid and in there the same grid square to ensure sample homogeneity.

### Data Analysis

The general scheme for analyzing paraffin data is shown in **Supplementary Figure 7**. First, motion correction with MotionCorr2^36^ and CTF correction with CTFFind3,^37^ were applied to the recorded data, using the software FOCUS.^38^ Power spectra of the image were computed in a python program, and the intensities of the 6 spots for the (1,1) and (2,0) reflections were measured. For background subtraction, the intensity of a 50-pixel diameter box around the centre of each spot was integrated and subtracted by the integrated intensity of pixels in an adjacent box of the same diameter. The average of the background-subtracted intensities of all spots (marked as 4 Å resolution spots) was normalized against the background-subtracted intensity from the first image. Measurements from 10 paraffin crystals were taken to ensure reproducibility of the results. A single exponential was fitted to each measurement, and *N*_*e*_ and values were computed for each measurement.

For data analysis of purple membrane images (**Supplementary Figure 9**), images were motion-corrected and CTF-corrected as above. After this, the images were unbent using the MRC software system integrated into the FOCUS software, which is a procedure that increases the SNR of the computed diffraction spots. The computed Fourier transform of purple membrane images showed many reflection spots across different resolutions. Thus, to track the radiation damage in different resolution zones, the reciprocal space was divided into different resolution zones as follows: 6 Å: 6-7 Å, 8 Å: 7-9 Å, 10 Å: 9-12 Å, 20 Å: 12-35 Å. The cutoff at 6 Å was chosen as the SNR of reflection spots at higher resolution was too low for a statistically significant analysis. The mean of background-corrected amplitudes in different resolution zones was squared and plotted against the accumulated electron doses. A single exponential was fitted to each measurement, and *N*_*e*_ values were computed for each measurement.

For data analysis of TMV images (**Supplementary Figure 11**), images were first motion-corrected and CTF-corrected as above. Then, power spectra of recorded micrographs were computed. These were background-subtracted, using a radial average profile. The TMV peaks at 23 Å were then identified, and their intensities were integrated. Peak intensities were normalized with the peak intensities from the first image, and the result was plotted against the accumulated electron dose. Again, a single exponential was fitted to each measurement, and *N*_*e*_ values were computed for each measurement. For per-frame B-factors calculation, the dataset was analyzed in Relion 5. Micrographs were motion corrected using Relion’s own implementation and then processed using the standard Relion processing pipeline.

## Code availability

Python codes used for evaluation of the paraffin crystal data are available at https://github.com/LBEM-CH/paraffin-analyzer

## Reporting Summary

(Will be provided.)

## Data Availability

The data sets generated in this work are available at the BioImage Image Archive (EMBL-EBI), accession code S-BIAD2466 (DOI: 10.6019/S-BIAD2466).

## Acknowledgements

We thank Richard Henderson for providing a first batch of BR samples, Bob Glaser for providing the protocol to prepare paraffin grids, Massimo Kube for helping in data processing in Relion, and Daniel Mann and Carsten Sachse for providing TMV samples. We acknowledge support by Thermo Fisher Scientific from Erik Kieft and Mark van Rijt for initial trainings for operating the instrument. V.K. acknowledges Nisika for useful discussions during the preparation of the manuscript.

This work was in part supported by the Swiss National Science Foundation (grant 200021_200628), and by the European Union (ERC 4D-BioSTEM, No 101118656). Views and opinions expressed are, however, those of the authors only and do not necessarily reflect those of the European Union or the European Research Council Executive Agency. Neither the European Union nor the granting authority can be held responsible for them.

## Author contributions

V.K. performed all experiments, optimized the RF Cavity, analyzed the data and wrote the manuscript. J.R. assisted in optimizing the RF cavity and maintained the microscope. C.K.V. froze the TMV grids. I.M. and R.C.G. assisted in optimizing the RF Cavity system. D.H. and D.F. prepared BR samples. H.S. and J.R. supervised the project. H.S. conceived the project. All authors contributed to the manuscript.

## Additional Information

**Supplementary Information** accompanies this paper at https:….

