## Supplementary Information for "Pulsed-electron illumination does not reduce beam damage for imaging biological macromolecules"

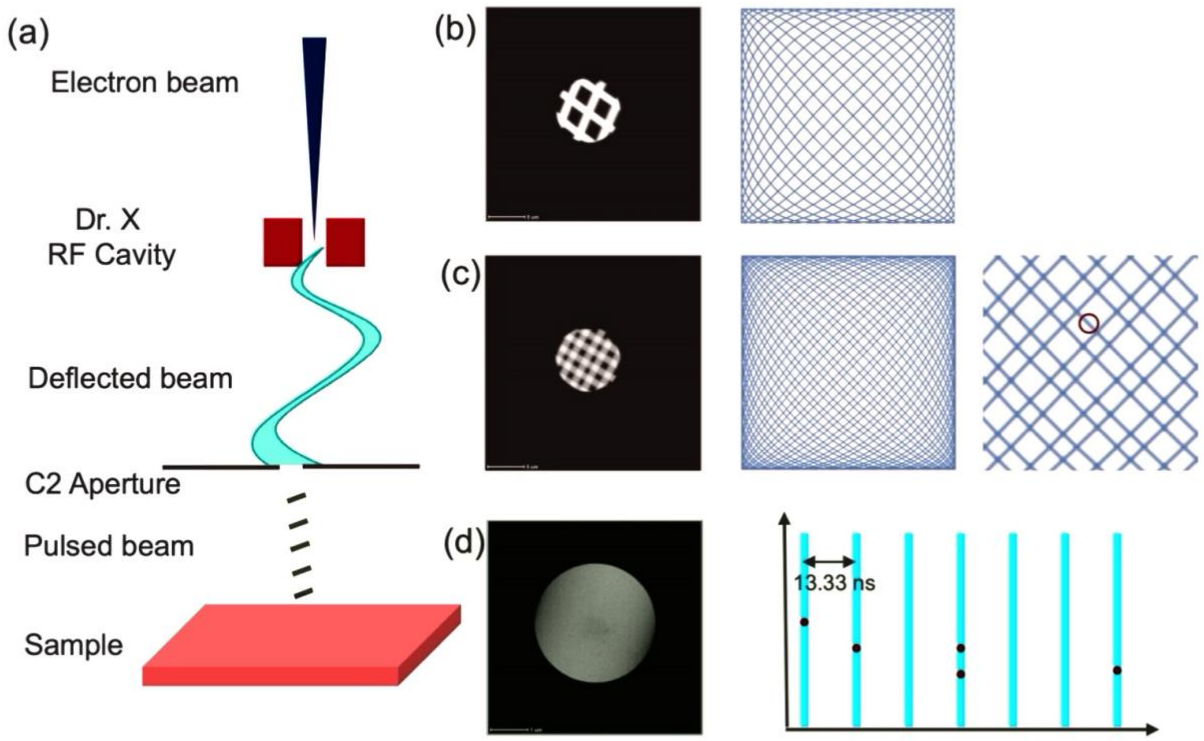

**Supplementary Figure 1. a** Schematic for the working of RF cavity. Experimental electron beam image and simulated Lissajous patterns **b** with and **c** without phase shift showing 32×33 crossings. **d** Finally, the chopped electron beam with corresponding temporal profile, showing spacing of 13.33 ns between each pulse corresponding to difference between two RF modes (75 MHz). First the electron beam is focused onto the RF cavity, where a strong magnetic field deflect the electron-beam to form a Lissajous pattern. After that, a small phase shift is applied between the two RF modes, to separate two beams riding on top of each other. Then, using the combination of condenser aperture, spot size, illumination and magnification, the electron beam is chopped, while a homogenous electron beam covers the full aperture. Now, as the beam is making a full circle taking 13.33 ns and only a very small section of beam is able to go through the aperture, a pulsed-electron beam with temporal spacing of 13.33 ns is created.

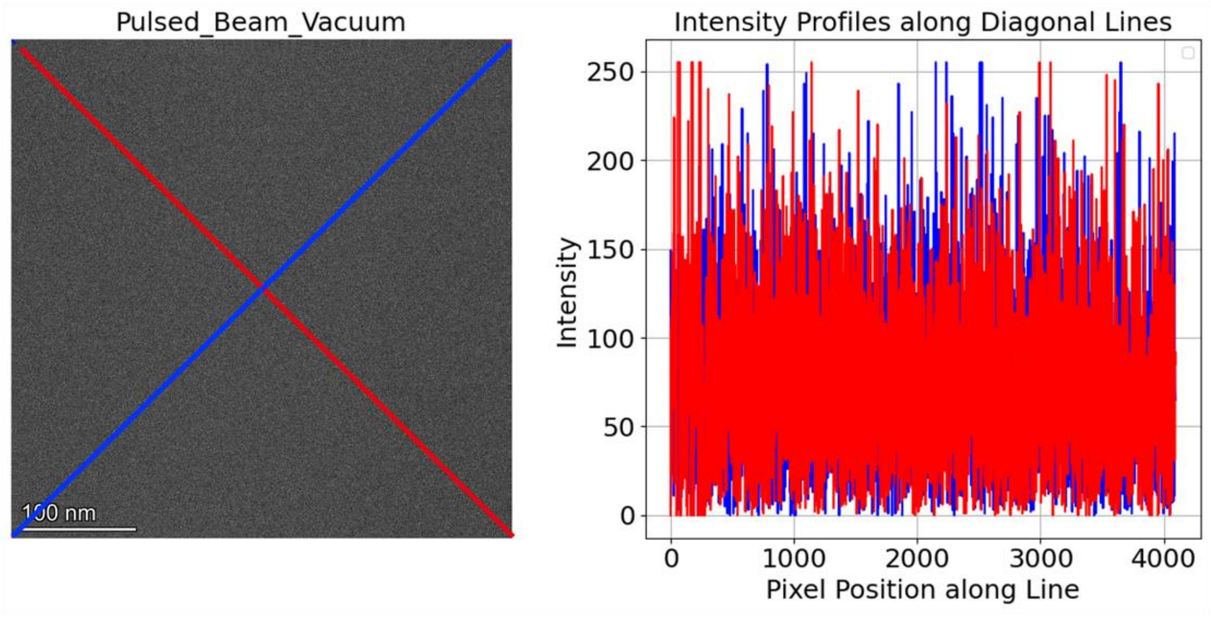

**Supplementary Figure 2.** TEM image over vacuum and corresponding intensity profiles along the diagonals demonstrating pulsed-beam beam homogeneity. Exposure time is 2 seconds.

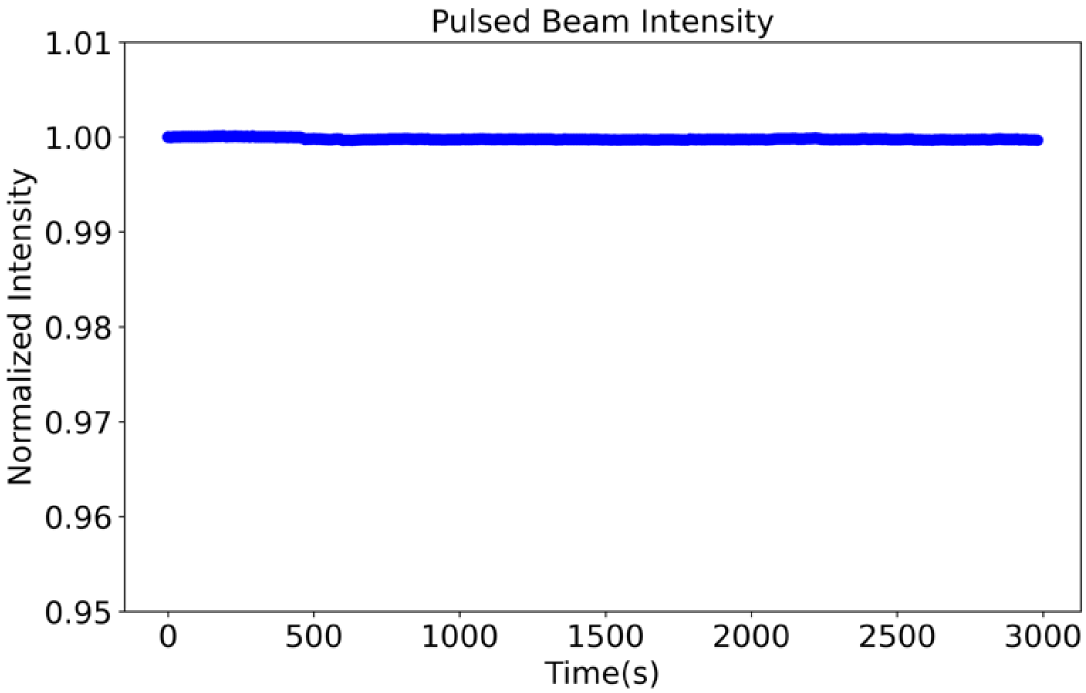

**Supplementary Figure 3.** Normalized intensity of the pulsed beam over time, showing a stable beam for 50 minutes, which was found suitable for imaging.

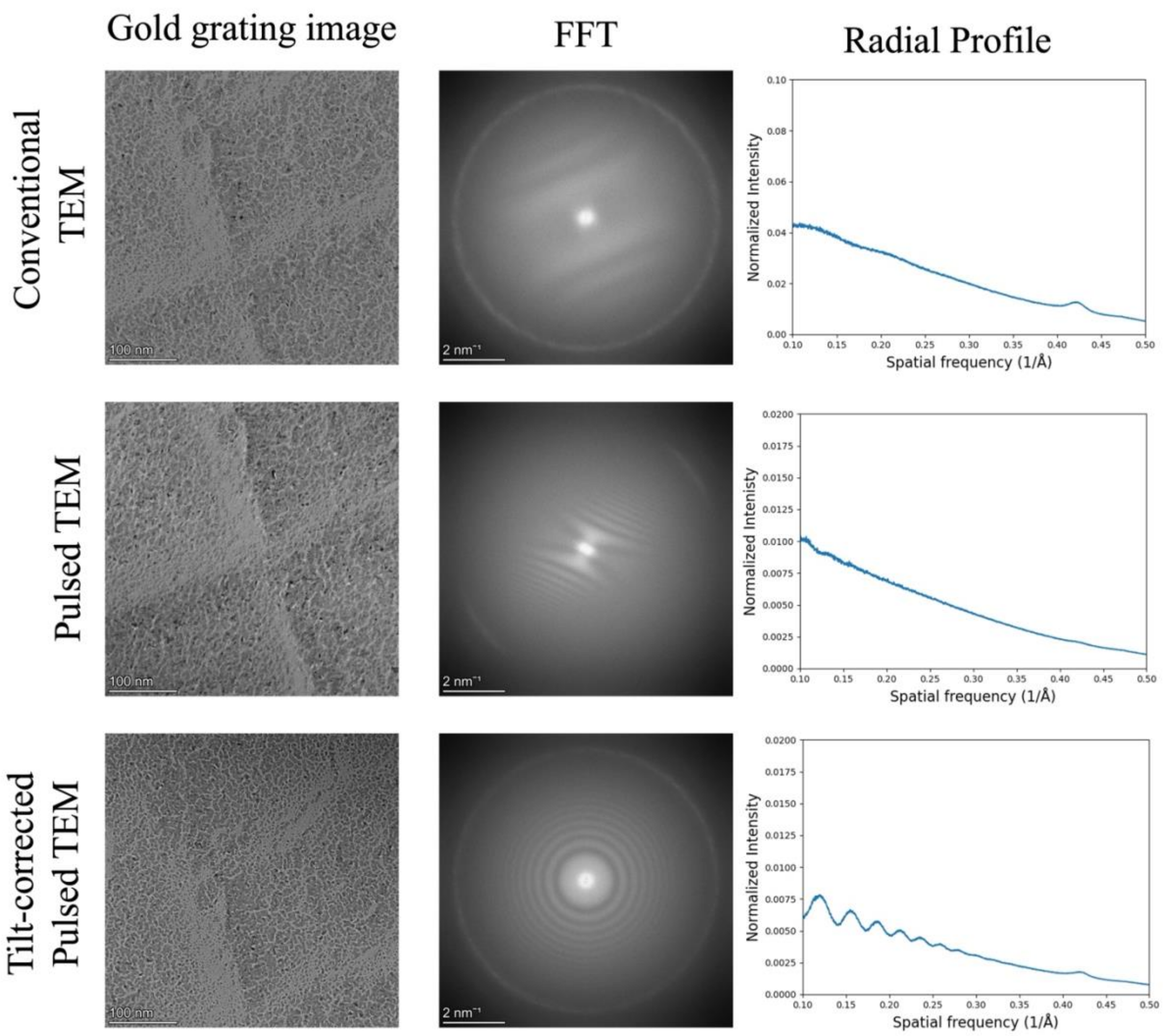

**Supplementary Figure 4.** TEM images of gold grating with different electron illumination and corresponding Fourier transform of images, showing information transfer in different directions up to the Nyquist limit (Magnification: 75kx, Pixel size: 1.05 Å). The DrX.Works cavity induces a strong beam tilt, leading to partial information transfer (central row), which is corrected by introducing tilt in the gun (bottom row). The radial profile shows a similar performance with conventional (top) and with tilt-corrected pulsed (bottom) operation. The conventional and the tilt-corrected pulsed TEM profiles show a similar gold diffraction intensity at 0.425 $Å^{-1}$ (2.35 Å).

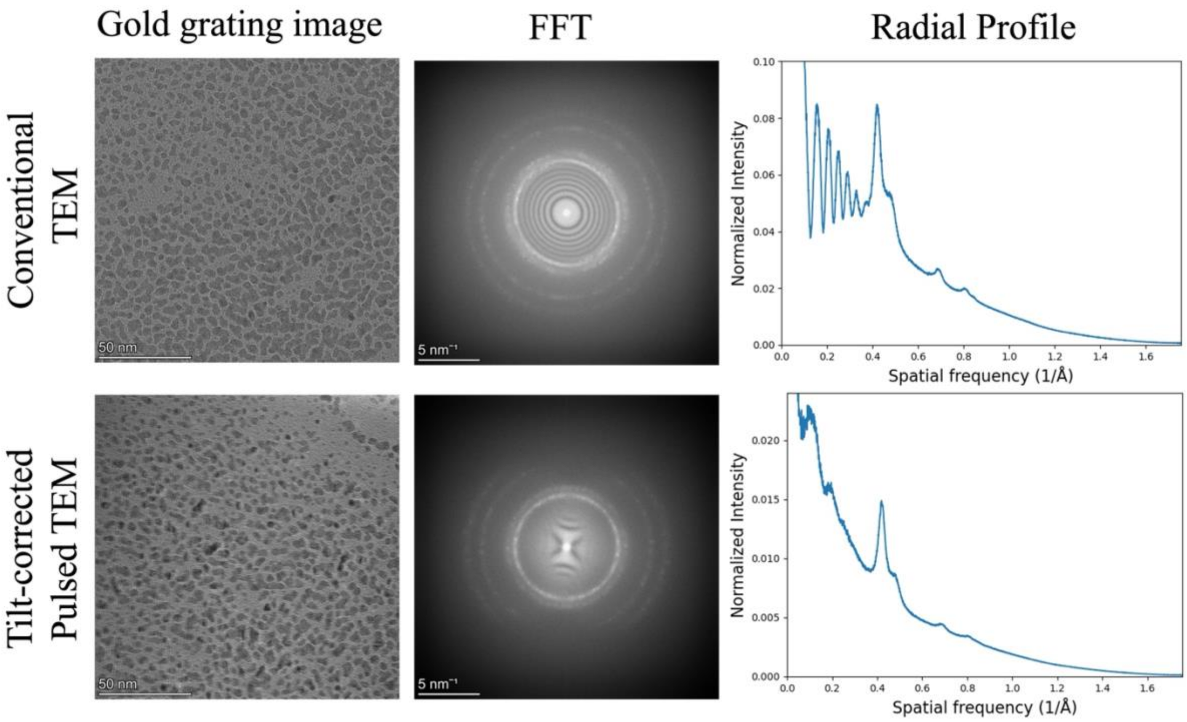

**Supplementary Figure 5.** High-resolution TEM images of gold grating with different electron illumination (left), and corresponding Fourier transforms of the images (middle), showing information transfer in different directions up to the Nyquist limit (Magnification: 195kx, Pixel size: 0.403 Å). Radial profiles (right) show for both, pulsed and conventional TEM illumination, information up to very high frequencies. The Image acquired with the pulsed mode shows partial loss of information in higher order diffraction rings 0.687 $Å^{-1}$ (1.45 Å), and 8.07 $Å^{-1}$ (1.24 Å), resulting from combination of increased beam tilt and loss of spatial coherence. The data for the pulsed mode were taken at lower defocus, resulting in lower intensity values for pulsed illumination.

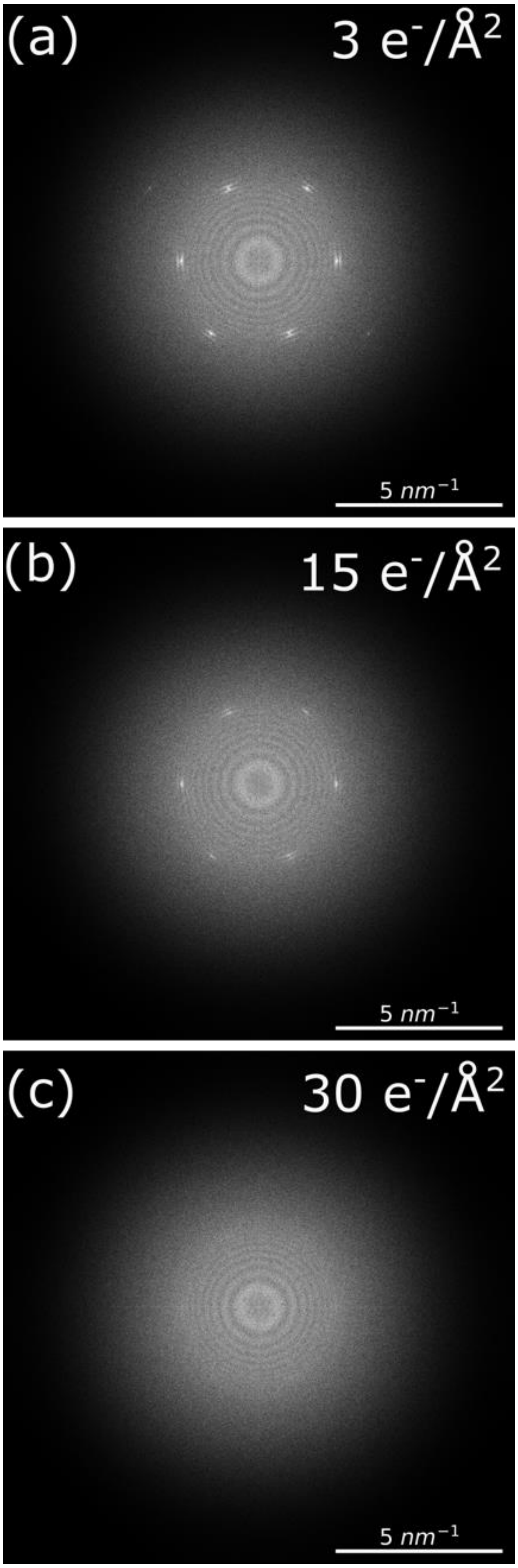

**Supplementary Figure 6.** Representative computed power spectra showing fading of paraffin diffraction spots with accumulated electron dose. **a** Power spectrum of the first recorded image. **b** Power spectrum of an image after 12 $e^-/Å^2$ had been given to the sample before. **c** Power spectrum of an image after 27 $e^-/Å^2$ had been given to the sample before.

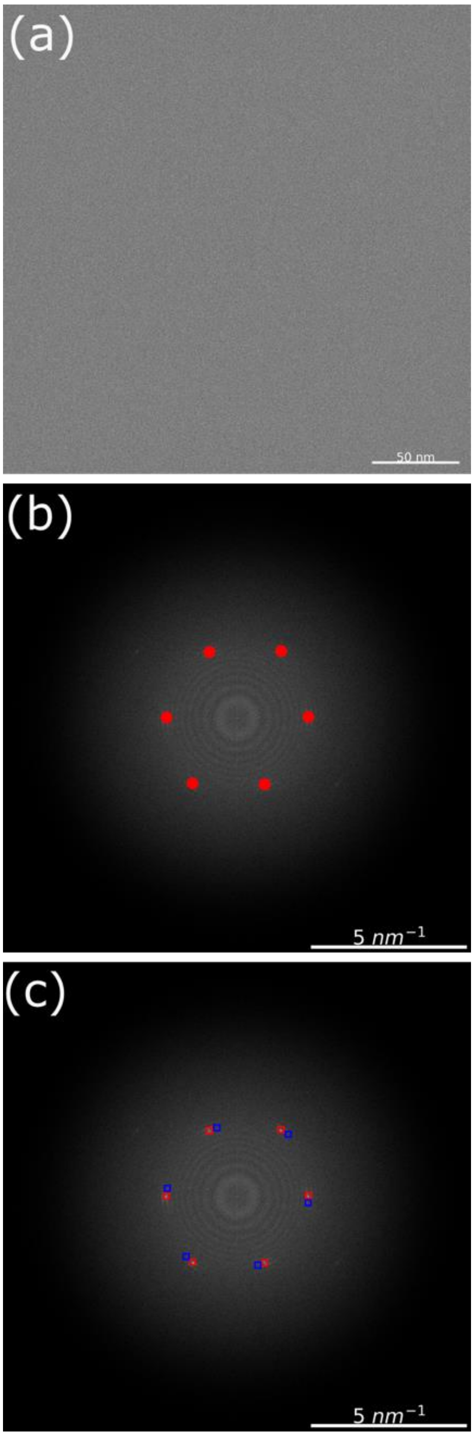

**Supplementary Figure 7.** Data processing pipeline for paraffin data (for both random and pulsed illumination). **a** First, the image was drift corrected using MotionCorr2 and CTF corrected (phase flipping). **b** Then, from the computed power spectrum of the image, the six innermost diffraction peaks ((110) and (200) reflections and their Friedel pairs) were identified. **c** For each peak, the intensity was integrated within a box of 50*50-pixels centered on the peak. For background subtraction, the intensity within a box of the same size but placed adjacent to the peaks was integrated, and this value was subtracted from the peak intensity. The mean of the background-subtracted intensities was then used for plotting against the cumulative electron dose.

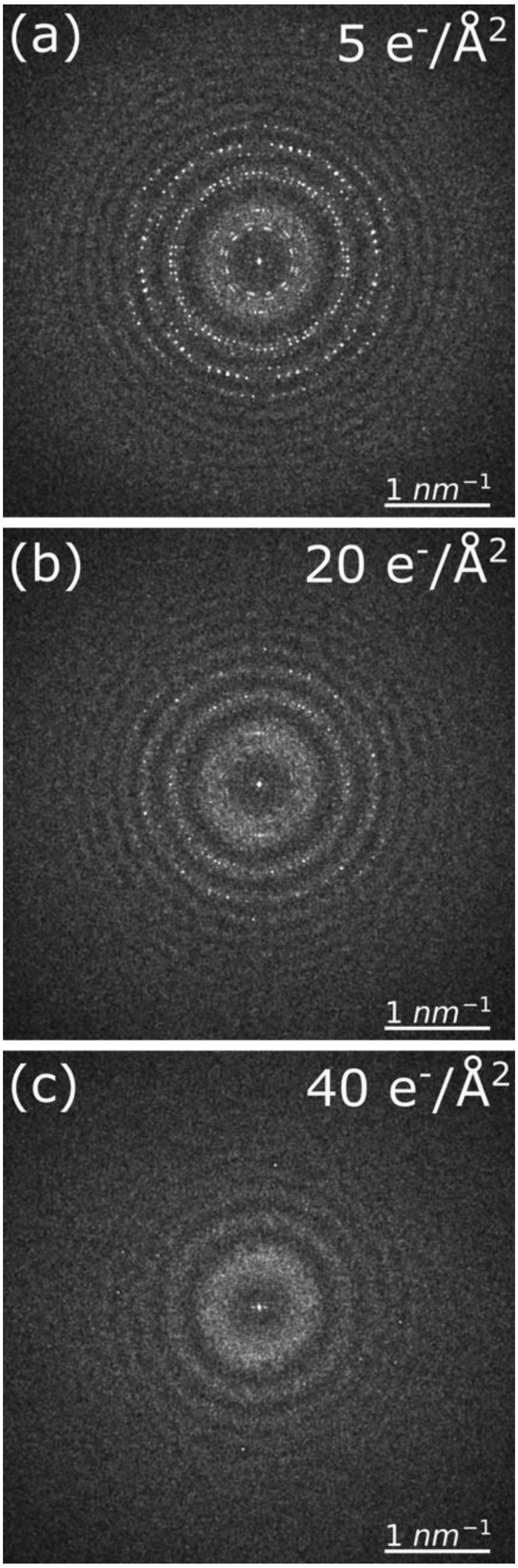

**Supplementary Figure 8.** Representative power spectrum showing fading of purple membrane diffraction spots with accumulated electron dose. The computed power spectra were Fourier-cropped from 4096*4096 to 1024*1024 pixels for better visualization of diffracted spots. **a** The power spectrum of the first recorded image. **b** Power spectrum after prior 15 $e^{-}/Å^2$. **c** Power spectrum after prior 35 $e^{-}/Å^2$.

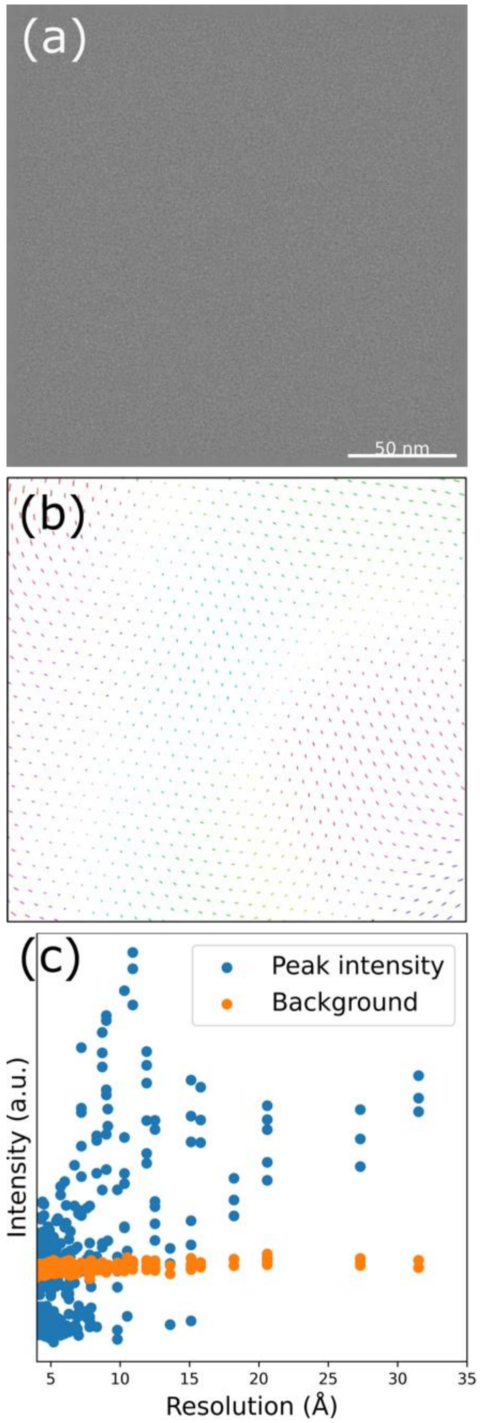

**Supplementary Figure 9.** Data processing pipeline for purple membrane crystal images. **a** First the image was drift corrected and CTF corrected. **b** the image was then unbent to compensate for protein crystal distortions, using the FOCUS software to enhance the intensity of the diffracted spots. The lattice unbending vectors shown here have a 10x exaggerated length for better visualization. From the FFT of the image, background-corrected peak intensities were then noted using MMBOXA script from the MRC software package, implemented in FOCUS. **c** Background-subtracted peak intensities and the background intensity surrounding each peak are plotted as a function of the peak resolution. From this, peaks were categorized into four different resolution zones (6 Å: 6-7 Å, 8 Å: 7-9 Å, 10 Å: 9-12 Å, 20 Å: 12-35 Å) for further analysis.

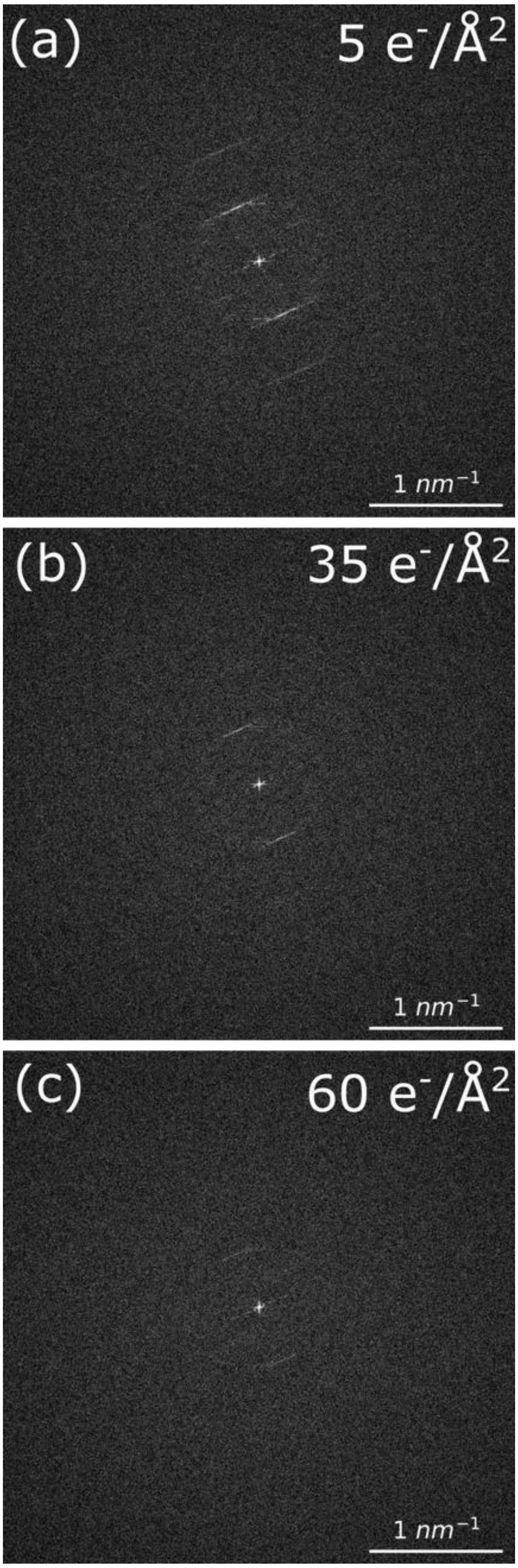

**Supplementary Figure 10.** Representative power spectra showing fading of diffracted rings with accumulated electron dose for TMV. The power spectra were Fourier cropped from 4096*4096 to 1024*1024 pixels for better visualization of diffracted rings. **a** Power spectrum of the first recorded image. **b** Power spectrum after prior 30 $e^-/Å^2$. **c** Power spectrum after 55 prior $e^-/Å^2$.

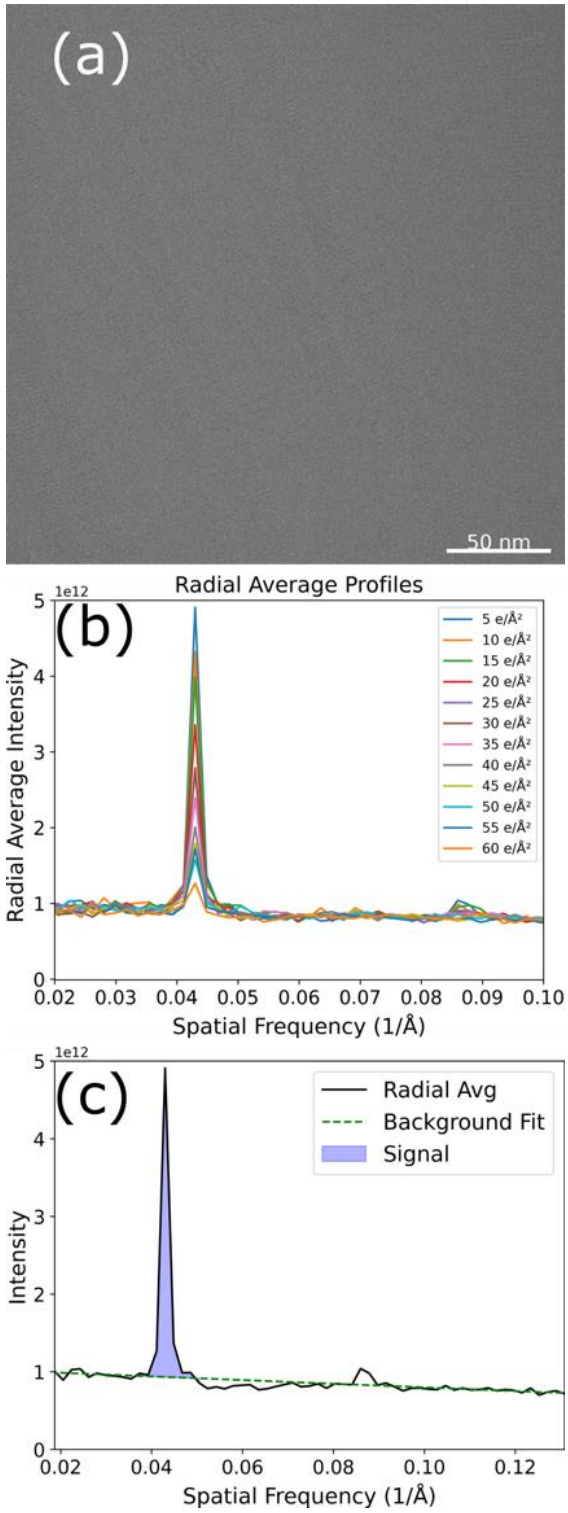

**Supplementary Figure 11.** Data processing pipeline for images of TMV embedded in vitreous ice. **a** First, images were drift corrected and CTF corrected as in previous datasets. **b** Then, from the computed power spectrum of the image, the radial average of the intensity was calculated, using a binning factor of 5 pixels. The first and second reflections at 23 and 11.5 Å resolution are discernible. **c** the background subtraction was performed and the integrated area under the first peak was measured and plotted against accumulative electron dose for 23 Å resolution peak.

**TMV Single-particle Analysis (SPA)**

Acquiring large datasets of TMV for a single particle analysis was challenging due to the requirement of regularly flashing the coldFEG tip (which normalizes electron optics), the inability to change magnification to large fields of view (required in automated data acquisition for performing Eucentric height adjustments and finding ice holes, etc.), and the inability to adjust the defocus value by larger amounts as this would have introduced excessive beam tilt.

Nevertheless, we have acquired a small dataset of TMV for SPA reconstruction, which was processed in Relion5. Data were acquired at an accelerating voltage of 300 kV with physical pixel size of 0.6475 Å at the detector level. Each image had a dose of 50 $e^-/Å^2$, which was processed in Relion as 50 frames. Thus, for per-frame B-factor calculation, each frame has a dose of 1 $e^-/Å^2$. The results have been summarized below:

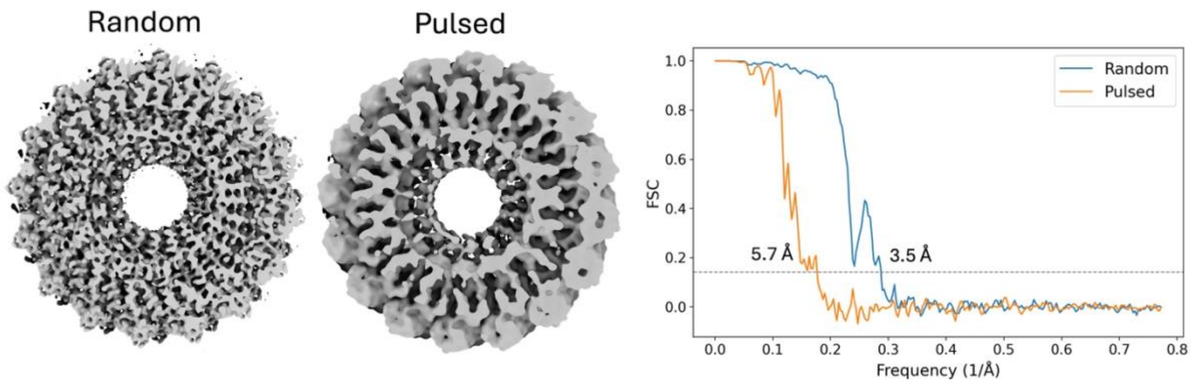

**Supplementary Figure 12.** Cross-sectional view of TMV maps for Random and Pulsed illumination and corresponding FSC plots for the TMV 3D reconstruction, comparing random with pulsed illumination. The lower resolution achieved with pulsed illumination may be due to beam stability and beam tilt issues.

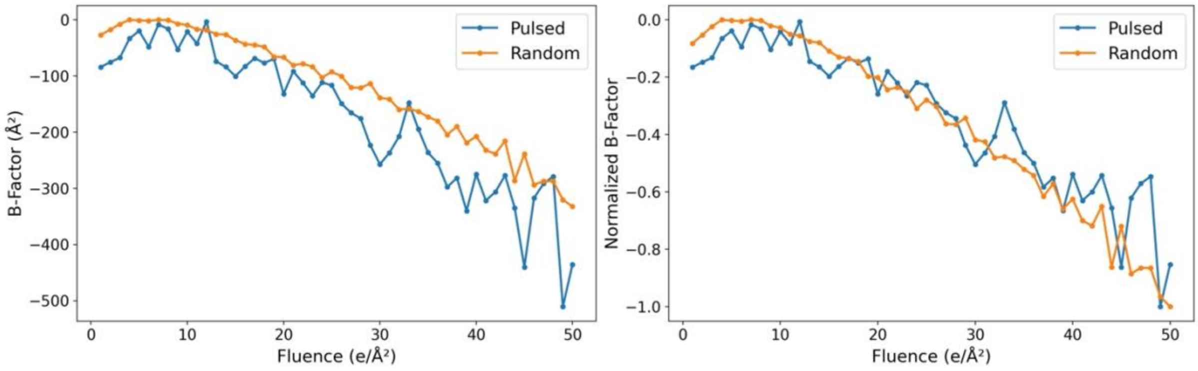

**Supplementary Figure 13. a** Per-frame B-factors, and **b** normalized per-frame B-factors for TMV datasets, analyzed in Relion5 for both, pulsed and random electron beam illumination. B-factor values are lower and noisier for the pulsed illumination dataset, as the map resolution is lower (5.7 Å) compared to that recorded with random illumination (3.5 Å). However, the decay profiles that inform about beam damage accumulation, are similar, especially after normalization, again indicating that damage dynamics are similar for pulsed and random illumination modes.

**Supplementary Table 1.** Beam statistics for imaging under pulsed and random electron-beam configuration.

| **Sample** | **Paraffin** | | **Purple Membrane** | | **TMV** | |
|---|---|---|---|---|---|---|
| | Pulsed | Random | Pulsed | Random | Pulsed | Random |
| **Pixel Size (Å)** | 0.65 | 0.65 | 0.52 | 0.52 | 0.65 | 0.65 |
| **Beam-Current (pA)** | 4 | 4 | 3 | 3 | 3 | 3 |
| **Beam Diameter (nm)** | 540 | 540 | 390 | 390 | 560 | 560 |
| **Dose-rate ($e^-$/Å$^2$ s)** | 1.35 | 1.36 | 2.08 | 2.10 | 1.10 | 1.10 |
| **Probability (1 $e^-$) per pulse** | 23.40 % | NA | 20.05 % | NA | 21.34 % | NA |
| **Probability (2 $e^-$) per pulse** | 3.78 % | NA | 2.61 % | NA | 3.02 % | NA |

**Supplementary Table 2.** Critical dose ($e^-/Å^2$) statistics for paraffin and TMV.

| **Measurement Number** | **Paraffin (4 Å)** | | **TMV (23 Å)** | |
|---|---|---|---|---|
| | **Pulsed** | **Random** | **Pulsed** | **Random** |
| 1. | 12.19 | 11.79 | 36.21 | 33.98 |
| 2. | 9.05 | 12.62 | 35.08 | 30.80 |
| 3. | 12.54 | 10.1 | 38.29 | 35.55 |
| 4. | 9.13 | 11.33 | 37.86 | 42.90 |
| 5. | 9.88 | 13.02 | 40.49 | 31.92 |
| 6. | 12.24 | 11.75 | 37.09 | 39.57 |
| 7. | 10.61 | 12.78 | 32.27 | 37.39 |
| 8. | 12.31 | 11.53 | 42.45 | 32.62 |
| 9. | 12.67 | 10.49 | 46.31 | 31.35 |
| 10. | 10.90 | 11.66 | 35.34 | 36.09 |
| **Mean ± Std** | 11.15 ± 1.47 | 11.71 ± 0.94 | 38.15 ± 3.84 | 35.22 ± 3.70 |

**Supplementary Table 3.** Critical dose ($e^{-}$/Å$^2$) statistics for purple membrane across different resolution zones.

| **Measurement Number** | **Purple Membrane** | | | | | | | |
|---|---|---|---|---|---|---|---|---|
| | **(6 Å)** | | **(8Å)** | | **(10 Å)** | | **(20 Å)** | |
| | **Pulsed** | **Random** | **Pulsed** | **Random** | **Pulsed** | **Random** | **Pulsed** | **Random** |
| 1. | 13.74 | 15.68 | 26.97 | 17.84 | 16.80 | 16.29 | 29.07 | 22.57 |
| 2. | 14.20 | 16.92 | 15.57 | 16.28 | 21.42 | 16.01 | 22.49 | 20.06 |
| 3. | 17.02 | 13.57 | 10.91 | 14.52 | 17.78 | 12.38 | 20.99 | 14.94 |
| 4. | 19.65 | 10.81 | 15.33 | 12.86 | 14.38 | 19.10 | 18.07 | 17.85 |
| 5. | 18.28 | 23.33 | 15.13 | 15.01 | 14.50 | 19.41 | 24.28 | 15.74 |
| 6. | 22.02 | 12.17 | 14.45 | 16.70 | 14.51 | 16.17 | 16.82 | 35.21 |
| 7. | 9.61 | 14.04 | 21.04 | 16.05 | 14.80 | 15.97 | 18.31 | 22.22 |
| 8. | 23.25 | 15.07 | 15.37 | 14.98 | 19.13 | 18.53 | 17.08 | 17.07 |
| 9. | 14.72 | 11.36 | 17.29 | 13.75 | 14.99 | 16.02 | 20.44 | 21.15 |
| 10. | 13.83 | 12.39 | 14.09 | 16.07 | 17.13 | 16.89 | 16.41 | 19.78 |
| **Mean ± Std** | 16.64 ± 4.01 | 14.63 ±3 .41 | 16.62 ± 4.21 | 15.41 ± 1.40 | 16.54 ± 2.25 | 16.68 ± 1.92 | 20.40 ± 3.80 | 20.66 ± 5.45 |
